# Multiplexed CRISPR gene editing in primary human islet cells with Cas9 ribonucleoprotein

**DOI:** 10.1101/2023.09.16.558090

**Authors:** Romina J. Bevacqua, Weichen Zhao, Emilio Merheb, Seung Hyun Kim, Alexander Marson, Anna L. Gloyn, Seung K. Kim

## Abstract

Successful genome editing in primary human islets could reveal features of the genetic regulatory landscape underlying β cell function and diabetes risk. Here, we describe a CRISPR-based strategy to interrogate functions of predicted regulatory DNA elements using electroporation of a complex of Cas9 ribonucleoprotein (Cas9 RNP) and guide RNAs into primary human islet cells. We successfully targeted coding regions including the *PDX1* exon 1, and non-coding DNA linked to diabetes susceptibility. CRISPR/Cas9 RNP approaches revealed genetic targets of regulation by DNA elements containing candidate diabetes risk SNPs, including an *in vivo* enhancer of the *MPHOSPH9* gene. CRISPR/Cas9 RNP multiplexed targeting of two *cis*-regulatory elements linked to diabetes risk in *PCSK1*, which encodes an endoprotease crucial for insulin processing, also demonstrated efficient simultaneous editing of *PCSK1* regulatory elements, resulting in impaired β cell *PCSK1* regulation and insulin secretion. Multiplex CRISPR/Cas9 RNP provides powerful approaches to investigate and elucidate human islet cell gene regulation in health and diabetes.

## INTRODUCTION

Impaired pancreatic islet function underlies nearly all forms of diabetes mellitus, including type 1 (T1D) and type 2 diabetes (T2D)^1^. Islets are comprised of clustered hormone-producing cells, called β, α, δ, ε and PP cells, that govern glucose and other key regulators of metabolism. Genetic and acquired risks are thought to impact islet function and promote diabetes development^2–4^. In both T1D and T2D, genetic risk has been linked to non-coding DNA variants, with a preponderance of these located in active islet gene regulatory regions called enhancers^5–6^. Recent studies show that islet enhancers change their chromatin accessibility during lineage progression and maturation^7^, upon glucose stimulation^8^, and upon cytokine exposure^9^. However, challenges in studying human islet enhancers, promoters and other *cis*-regulatory elements, have limited our understanding of how these are mechanistically linked to diabetes risk^10–11^.

Attempts to study gene regulation in primary human islet cells face multiple hurdles including: (1) the relatively small number of islet cells per pancreas, (2) their lack of expansion - islet cells are largely non-dividing, (3) the characteristic clustering of islet cells that limits efficient genetic targeting, (4) the harsh nucleolytic environment of the exocrine pancreas surrounding native islets, and (5) a lack of methods to target or edit non-coding DNA elements in primary islets. To address these technical gaps, we innovated methods for genetic modification of primary human islet cells using CRISPR/Cas9-based approaches. We previously showed efficient gene editing of primary human islet cells using lentiviruses simultaneously encoding Cas9, a sgRNA and GFP^12^. This system, combined with our transient dispersion of primary islet cells followed by reaggregation into organoids called ’pseudoislets’ (reviewed in^13^), allowed efficient gene editing of both coding and non-coding genomic regions in primary human islet cells. However, lentivirus-based CRISPR/Cas9 targeting is relatively labor-intensive, requiring cloning of candidate single-stranded guide RNAs (sgRNAs), followed by production of virus in sufficient titers, then isolation of infected cells for analysis. These multiple steps reduce yields for cell or nucleic acid-based assays, and limit ease of scalability or genetic screens.

Electroporation of Cas9 ribonucleoprotein pre-loaded with sgRNAs (CRISPR/Cas9 RNP) has been successfully used for gene editing in challenging cellular targets, including embryonic stem cells, induced pluripotent stem cells, tissue stem cells and T lymphocytes^14–17^, but has not previously been reported for adult human islet cells, which are post-mitotic. Direct delivery of CRISPR/Cas9 RNP complexes can bypass the requirement for transcription and translation, allowing rapid and efficient genome editing. In contrast to lentiCRISPR approaches, the CRISPR/Cas9 RNP does not integrate into the genome. Thus, the only genomic modification introduced is the specific gene edit. The short half-life of the CRISPR/Cas9 RNP complex reduces off-target effects, and the chance for insertional mutagenesis, and immune responses^18–19^. The simplicity of this approach can also enable simultaneous (’multiplex’) editing of multiple genetic targets^16^ like promoters and enhancers. Here, we investigated use of CRISPR/Cas9 RNP-based multiplex targeting of *cis*-regulatory elements in primary human islet cells.

## RESULTS

### CRISPR/Cas9 RNP electroporation to target PDX1 in primary human islet cells

To innovate CRISPR gene editing in primary human islet cells, we electroporated Cas9 protein fused with GFP with sgRNAs into dispersed human islet cells (**Fig 1A**: Methods). Afterwards, islet cells reaggregated to form pseudoislets that were assayed after 6 days, as we previously reported using the lentiCRISPR approach^12^.

**Figure 1.**
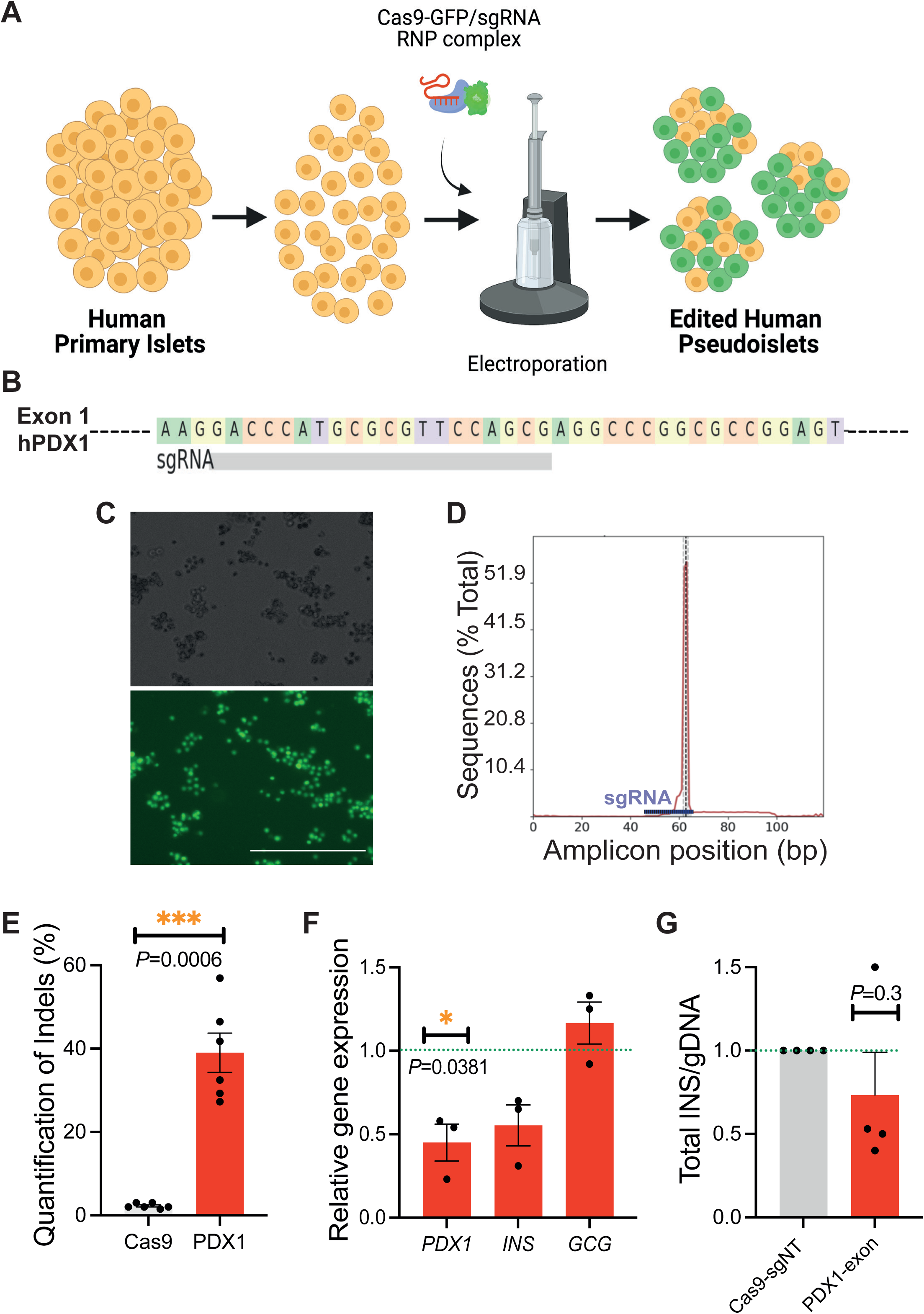
Efficient CRISPR/Cas9 RNP mediated targeting of PDX1 in primary human islet cells. (A) Schematic of the human pseudoislet electroporation system with CRISPR/Cas9 RNP complexes. (B) Fragment of human *PDX1* exon 1 sequence, showing the sgRNA sequence in grey. (C) Human pseudoislets 1 day post-electroporation with Cas9-EGFP RNP and PDX1 sgRNAs complexes (top panel: bright field; bottom panel: blue light, 488 nm, scale bar: 500 μm). (D) Quantification of editing frequency mapped to the reference amplicon using CRISPResso analysis. As expected, the mutations cluster around the predicted cleavage position based on the sgRNA sequence (expected cleavage site indicated by a vertical dotted line; sgRNA sequence: violet line). (E) Quantification of indels after CRISPR/Cas9 RNP targeting of PDX1 (PDX1, red) or using a control sgRNA sequence (Cas9, grey) (n = 6 independent human islet donors). (F) qRT-PCR of pseudoislets, CRISPR-PDX1 (red), normalized to the CRISPR-Control (n = 3 independent donors). (G) Total insulin content of CRISPR/Cas9 electroporated cells normalized to genomic DNA (gDNA) content (n = 4 independent donor samples). Data are presented as mean values ± SE. Two-tailed t tests were used to generate *P* values. See also Figure S1.

To optimize CRISPR/Cas9 RNP electroporation, we systematically varied voltage and concentration of Cas9 RNP ’preloaded’ with sgRNA specific for exon 1 of *PDX1* (**Fig 1B**; **Fig S1A**, Methods), then visualized Cas9-GFP fluorescence one day after electroporation (**Fig 1C**). To assess toxicity of the system, we evaluated viability following electroporation with CRISPR/Cas9 RNP and 2 control sgRNAs as compared to non-electroporated pseudoislets 5 days following electroporation, and detected an average of 70% live cells (**Fig S1B-E**, Methods). Following 6 days of *in vitro* pseudoislet culture, we extracted genomic DNA and performed next-generation sequencing of the indexed amplicons (Methods). Insertion-deletion mutations (indels) were detected in an average of 40% of sequences by CRISPResso analysis^20^ (**Fig 1D-E**; **Fig S1F**). After CRISPR/Cas9 targeting of *PDX1*, qRT-PCR revealed ∼60% reduction of mRNA encoding PDX1 (**Fig 1F**). In addition, we observed a significant reduction in total INSULIN protein levels, similarly to our prior findings using lentiviral-based CRISPR (**Fig 1G**: ^12^). These results motivated studies to explore the potential of this CRISPR/Cas9 RNP approach to target non-coding DNA elements in primary human islet cells.

### Targeting of candidate *cis*-regulatory genomic regions in human islet cells

We have used lenti-CRISPR to identify native human β cell enhancer elements regulating *SIX2* and *SIX3*^12^, but it is unknown whether CRISPR/Cas9 RNP strategies can be used similarly. To assess this, we adapted CRISPR/Cas9 RNP-based methods to induce a deletion in a putative cis-regulatory element in human islets and to identify the effector transcripts. We investigated a putative regulatory element in the intronic region of *PITPNM2* linked by chromatin looping studies to *MPHOSPH9, PITPNM2* and *C12orf65*^8^ (**Fig 2A**; **Fig S2**). We electroporated dispersed islet cells with CRISPR/Cas9 RNP complexes with two sgRNAs (sg1 and sg2) to target DNA encompassing this postulated regulatory element (**Fig 2A**), and visualized Cas9-GFP fluorescence one day after electroporation (**Fig 2B**).

**Figure 2.**
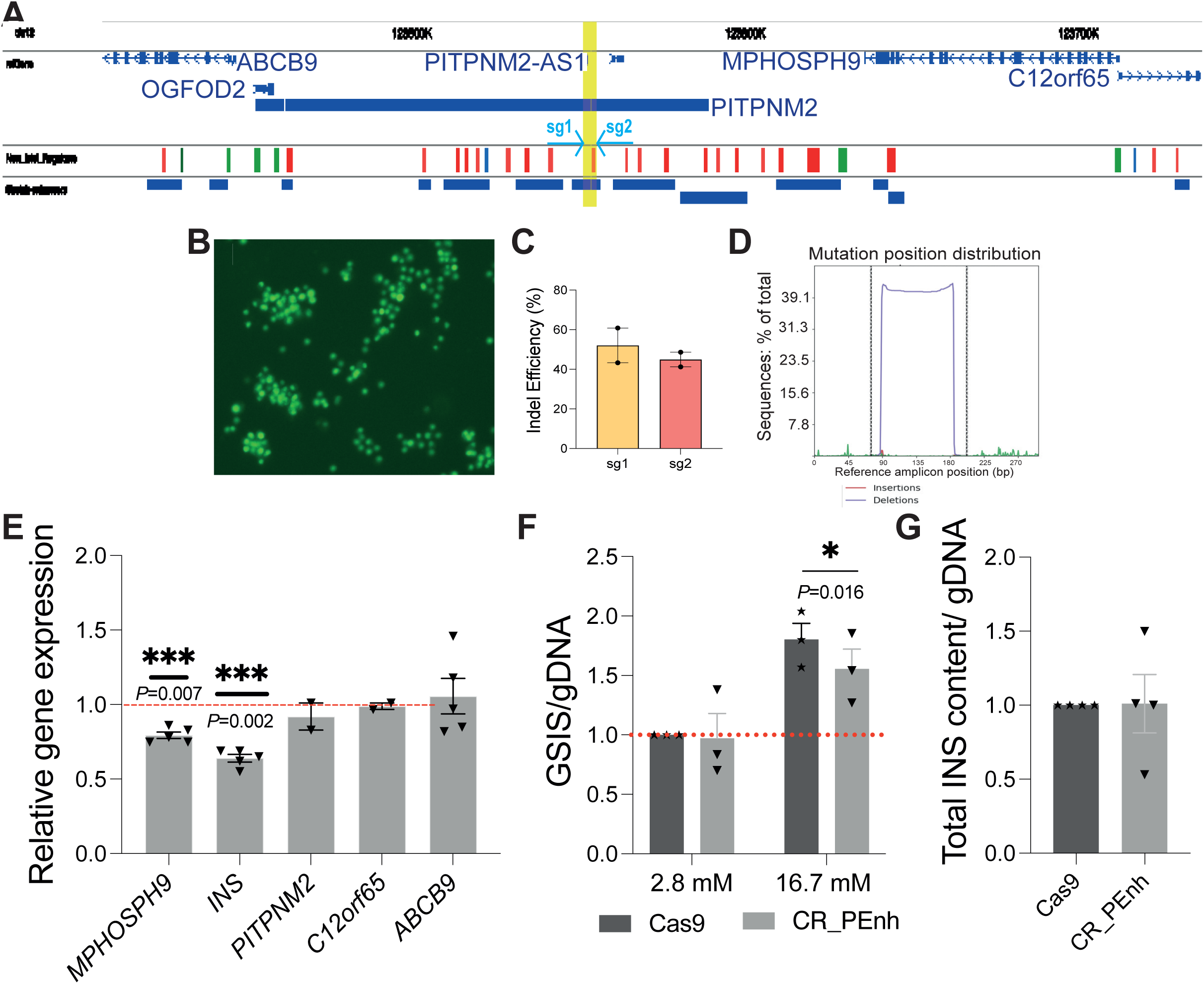
Linking candidate cis-regulatory genomic regions to their target genes in human islet cells. (A) Genome browser tracks of the *MPHOSPH9* loci, highlighting in yellow a candidate cis-regulatory genomic region within the *PITPNM2* gene, and linked by pC-HiC to *PITPNM2*, *MPHOSPH9* and *C12orf16*. See also Fig S2. The predicted enhancer (red, yellow highlight) was targeted with two sgRNAs (sg1 and sg2, turquoise arrows) designed to delete this element. (B) Human islet cells expressing GFP one day following electroporation with the CRISPR/Cas9 RNP complexes. (C) Quantification of editing frequency by each sg RNA mapped to the reference amplicon using CRISPResso analysis. See also Fig S3. (D) Histogram showing deletion of the region in between the two sgRNAs, generated with CRISPResso. (E) RT-qPCR showing reduced expression of *MSPHOSPH9* and *INS*, but not *PITPNM2*, *c12orf65* and *ABCC9* after deletion of the putative enhancer site with CR_PEnh compared to the Cas9 control. (F) Glucose-Stimulated Insulin secretion for the control (Cas9) versus CR_PEnh groups; (G) Total insulin content for the Cas9 control versus CR_PEnh groups. Data are presented as mean values ± SE. Two-tailed t tests were used to generate *P* values. \*\**P* < 0.05, \*\*\**P* < 0.01.

Controls included CRISPR/Cas9 RNP complexes with a non-targeting sgRNA. Following reaggregation and pseudoislet culture, deep sequencing and CRISPResso analysis^20^ of the indexed targeted amplicons detected indels and deletion efficiencies of >40% (**Fig 2C-D**). Sequencing analysis of the likeliest genomic off-target sites (Methods) revealed that indels were undetectable in 5/5 of these sites (**Fig S3**). qRT-PCR revealed a significant reduction of *MPHOSPH9* mRNA while mRNA encoding PITPNM2, C12orf65 or the nearby *ABCB9* gene were unaltered (**Fig 2E**).

Unexpectedly, *INS* mRNA levels were also reduced after CRISPR/Cas9 RNP targeting (**Fig 2E**). While total insulin content did not change, we observed impaired glucose-stimulated insulin secretion after CRISPR/Cas9 RNP targeting (**Fig 2F-G**). Thus, our studies provide *in vivo* evidence that this enhancer is active in native β-cells, and that its activity impacts the expression of *MPHOSPH9* but not of other neighboring genes. Moreover, reduced *MPHOSPH9* expression was linked to impaired *INS* expression and glucose-stimulated insulin secretion. *MPHOSPH9* encodes MPP9, a protein required for formation of primary cilia^21^, organelles known to govern hormone secretion by islet α and β cells^22,23^.

### Multiplexed CRISPR/Cas9 RNP targeting of regulatory regions in human islet cells

Next, we investigated use of CRISPR/Cas9 RNP for multiplex targeting of *cis*-regulatory elements in human islet cells. *PCSK1* encodes for Prohormone Convertase 1/3 (PC1/3), an endopeptidase regulating processing of Proinsulin (**Fig 3A**), and linked by prior GWAS studies to human diabetes and obesity risk^24–25^. To target the established promoter and enhancer regulatory elements of *PCSK1* in native β cells (**Fig S4A)**, we electroporated primary human islets with CRISPR/Cas9 RNP complexes encoding Cas9-GFP and two sgRNAs to induce a deletion of *PCSK1* promoter (*PCSK1*-Prom) or enhancer sequences (*PCSK1*-Enh; **Fig 3A-B**). Non-targeting sgRNAs complexed with Cas9-GFP served as controls. After electroporation and pseudoislet *in vitro* culture, we performed sequencing (Mi-seq) of the targeted *PCSK1*-Prom (**Fig 3D**) and *PCSK1*-Enh amplicons (**Fig 3E**) and used CRISPResso analysis to quantify the degree and specificity of gene editing. With *PCSK1*- Prom sgRNAs (sg1 and sg2), we observed >90% of *PCSK1* promoter sequences were modified; by contrast *PCSK1*-Enh sequences were not edited (**Fig 3D**). Similarly, >85% of enhancer sequences were modified using *PCSK1*-Enh1 sgRNAs (sg3 and sg4), while PC1/3 promoter sequences were not altered (**Fig 3E**). Potential genomic off-target sites were also sequenced (Methods), and no indels were detected in 6/6 of these sites (**Fig S6**). Thus, CRISPR/Cas9 RNP complexes were effective for independently targeting *PCSK1* promoter and enhancer elements.

**Figure 3.**
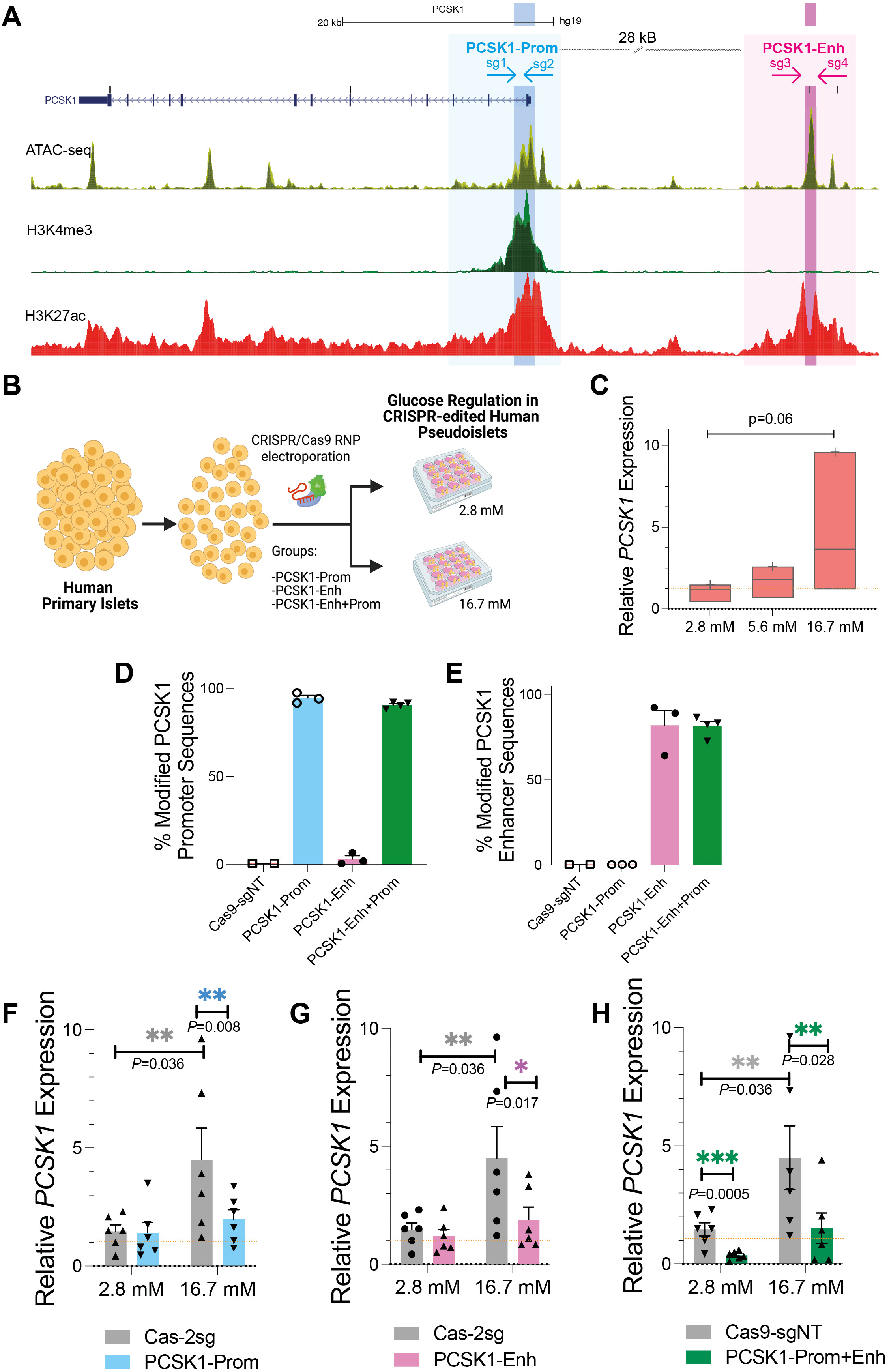
CRISPR/Cas9 RNP electroporation in human islet cells allows multiplex targeting of regulatory regions. (A) Genome browser tracks of *PCSK1* and its regulatory elements: *PCSK1* Promoter (*PCSK1*-Prom) and *PCSK1* Enhancer (*PCSK1*-Enh). Regulatory regions that show glucose-induced H3K27ac accessibility are highlighted: *PCSK1*-Prom (light blue, sg1 and sg2 are designed to induce a deletion of the turquoise region) and *PCSK1*-Enh (light pink, sg3 and sg4 are designed to induce a deletion of the dark pink region). Accessible chromatin regions in the human islets are shown by ATAC-seq, H3K4me3, and H3K27ac ChiP-seq. See also Fig S4. (B) Schematics of the human pseudoislet CRISPR/Cas9 electroporation approach used for targeting of *PCSK1*-Prom, *PCSK1*-Enh, or simultaneous targeting of *PCSK1*-Prom+Enh, followed by culture at either basal (2.8 mM) or high (16.7 mM) glucose concentrations. (C) qRT-PCR of *PCSK1* in pseudoislets at 2.8, 5.6 and 16.7 mM glucose (n = 3 independent donors). (D-E) CRISPResso quantification of editing efficiency on (D) *PCSK1* Promoter sequence and (E) *PCSK1* enhancer sequence, after targeting with CRISPR/Cas9 control (Cas9-sgNT), *PCSK1*- Prom, *PCSK1*-Enh or *PCSK1*-Enh+Prom (n = 3 independent donors for *PCSK1*-Enh, *PCSK1*-Prom and n = 4 for *PCSK1*-Enh+Prom. See also Fig S5, Fig S6 and Fig S7. (F-H) Measurements of *PCSK1* expression 5 days after CRISPR/Cas9 RNP targeting of (F) *PCSK1*- Prom, (G) *PCSK1*-Enh and (H) *PCSK1*-Enh+Prom compared to a control (Cas9-2sg) in human pseudoislets and culture at 2.8 mM versus 16.7 mM glucose (n = 5 independent donors). Data are presented as mean values ± SE. Two-tailed t tests were used to generate *P* values.

In human islets, combinatorial targeting of independent regulatory elements could allow molecular studies of polygenic diabetes risk. For dual targeting of *PCSK1*-Prom and PCSK1-Enh, we used sgRNAs to induce a deletion in both the Promoter and Enhancer elements (*PCSK1*-Prom+Enh; **Fig 3A-B**), separated by 28 kilobases. Both the *PCSK1*-Prom and *PCSK1*-Enh sequences were modified at similar levels (>80%) compared to targeting *PCSK1*-Prom or *PCSK1*-Enh regions alone (**Fig 3D-E**, **Fig S5 A-H**). Thus, CRISPR/Cas9 RNP targeting induced simultaneous deletions of two distinct *PCSK1* regulatory elements. We did not detect genomic off-target indels at any of the six potential sites after the dual targeting (Methods; **Fig S6**). We also performed indexed PCRs of three independent amplicons within the 28 kB region in between the targeted enhancer and promoter (internal amplicons 1, 2 and 3), as well as of a reference amplicon located outside the targeted region (**Fig S7A**). Following Miseq sequencing, we calculated the ratios of internal/reference amplicons, not detecting significant differences between the control and *PCSK1*-Prom+Enh (**Fig S7B**). We also measured int/Ref amplicon ratios for two of these amplicons and islets targeted with *PCSK1*-Prom and *PCSK1*-Enh (**Fig S7C-D**). As expected, the targeting of only one of the regulatory elements did not result in large deletions. Finally, we also performed PCR of the genomic region connecting enhancer and promoter, and confirmed that deletions spanning this region occurred in <20% of sequences after dual targeting (**Fig S8**).

*PCSK1* expression is regulated by glucose^8^, and we noted increased *PCSK1* mRNA levels in islet cells cultured in 16.7 mM glucose (’high’) compared to those in 2.8 mM glucose (’basal’; **Fig 3C**), corroborating prior reports. To assess the role of the *PCSK1*-Prom and *PCSK1*-Enh elements in glucose-dependent *PCSK1* expression, we targeted *PCSK1*-Prom, *PCSK1*-Enh, or *PCSK1*-Prom+Enh, then measured *PCSK1* mRNA levels in pseudoislets at 2.8 mM or 16.7 mM glucose (**Fig 3B,F-H**). As expected, cells electroporated with control Cas9- sgNT had increased *PCSK1* mRNA when cultured in high glucose compared to culture in basal glucose (**Fig 3F-H**). However, *PCSK1* mRNA induction was blunted after targeting *PCSK1*-Prom, *PCSK1*-Enh, or *PCSK1*-Prom+Enh (**Fig 3F-H**). Additionally, at low glucose concentration, *PCSK1* mRNA was reduced after targeting of *PCSK1*-Prom+Enh, but not in cells with *PCSK1*-Prom or *PCSK1*-Enh targeting alone (**Fig 3H**: *P<0.05*). These studies therefore reveal an additive requirement for both *PCSK1*-Prom and *PCSK1*-Enh in regulating *PCSK1* expression in basal glucose conditions.

We next evaluated roles of the *PCSK1*-Prom and *PCSK1*-Enh elements in glucose-dependent *PCSK1* expression in islet cells cultured in normoglycemic conditions (5.6 mM). In this case, we also observed reduced *PCSK1* mRNA targeting of *PCSK1*-Prom, *PCSK1*-Enh and *PCSK1*-Prom+Enh (**Fig 4A**). Levels of mRNAs encoding PCSK2 and GLUCAGON did not change following CRISPR targeting of *PCSK1* regulatory elements in cells cultured at 5.6 mM glucose concentration (**Fig 4A-B**), 2.8 mM or 16.7 mM glucose (**Fig S9C**). Glucose-dependent regulation of *INS* or *IAPP* (**Fig 4C-D** and **Fig S9A-B**) was also unaffected.

**Figure 4.**
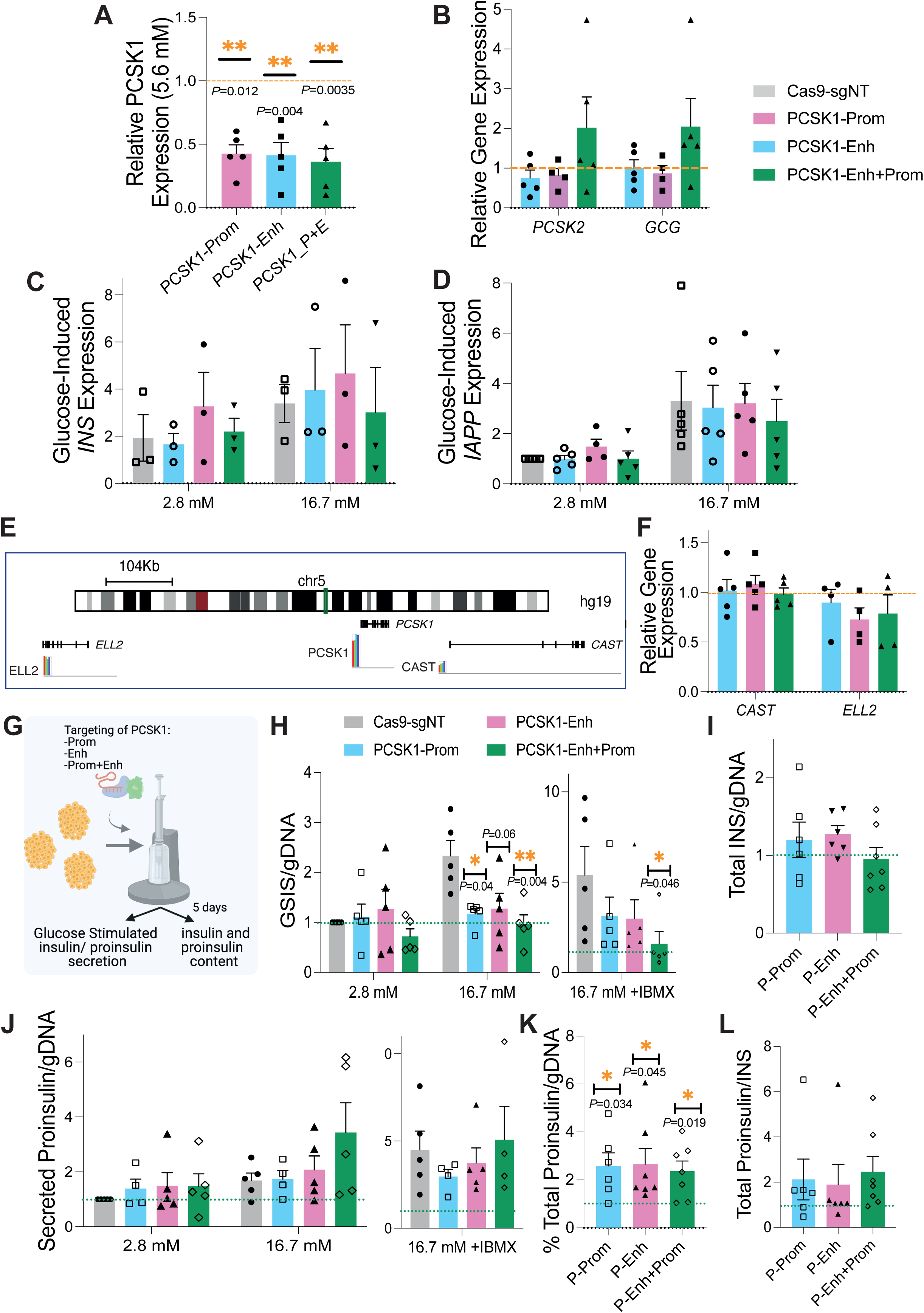
Selective impairment of *PCSK1* expression and impaired insulin processing and secretion following CRISPR/Cas9 RNP targeting of *PCSK1* regulatory elements. RT-qPCR after CRISPR/Cas9 RNP targeting of *PCSK1*-Prom (turquoise), *PCSK1*-Enh (pink) or *PCSK1*- Enh+Prom (green): (A-B) Following culture at 5.6 mM glucose, and measurement of mRNA levels of (A) *PCSK1* and (B) *PCSK2* and *GCG* (n = 5 donors). (C-D) Following culture at 2.8 mM versus 16.7 mM glucose and measurement of mRNA levels of glucose regulated (C) *INS* (n = 3) and (D) *IAPP* expression (n = 5). (E) Scheme of the *PCSK1* locus, showing *PCSK1* neighboring genes. (F) RT-qPCR of *CAST* and *ELL2* following CRISPR/Cas9 RNP targeting of *PCSK1* regulatory regions (n = 5 independent donors for *CAST* and n= 4 for *ELL2*). Data are presented as mean values ± SE. Two-tailed t tests were used to generate *P* values. \**P* < 0.05. (G) Scheme of electroporation of CRISPR/Cas9 RNP complexes with sgRNAs targeting *PCSK1*-Prom (turquoise bars), *PCSK1*-Enh (pink bars), *PCSK1*-Enh+Prom (green bar) or the control Cas9-sgNT followed by measurements of: (H) Glucose-stimulated Insulin Secretion (n = 5), (I) Total INS content (n = 5), (J) Glucose-stimulated Proinsulin Secretion (n = 4), (K) Total Proinsulin Content (n = 6-7), (L) Ratio of Proinsulin/Insulin (n = 6-7). See also Fig S8. Data are presented as mean values ± SE. Two-tailed t tests were used to generate *P* values. \**P* < 0.05, \*\**P* < 0.005.

Likewise, levels of mRNAs encoding *ELL2* and *CAST*, two genes neighboring *PCSK1* and located in the same topologically associated domain (TAD) as the *PCSK1-*Prom and *PCSK1-* Enh elements, were not altered (**Fig 4E-F**). Thus, we achieved CRISPR/Cas9 RNP based targeting of two non-coding regulatory regions in human β-cells, and our *in vivo* studies revealed a selective impact on the expression of *PCSK1*.

### Impaired insulin processing and secretion after targeting *PCSK1* regulatory elements

To assess the impact of targeting *PCSK1* regulatory elements on β cell function, we measured processing and secretion of Proinsulin and Insulin (**Fig 4G**). After CRISPR/Cas9 RNP targeting of *PCSK1*-Prom, *PCSK1*-Enh, or *PCSK1*-Prom+Enh we observed impaired Insulin secretion after stimulation with 16.7 mM glucose (**Fig 4H**). Moreover, challenge with 16.7 mM + the secretion potentiator IBMX revealed impaired Insulin secretion selectively after *PCSK1*-Prom+Enh targeting (**Fig 4H**), compared to targeting *PCSK1-*Prom or *PCSK1-*Enh alone. By contrast, Proinsulin secretion and total Insulin content in the *PCSK1*-Prom, *PCSK1*-Enh, or *PCSK1*-Prom+Enh groups was not significantly changed compared to controls (**Fig 4I-J**). However, total Proinsulin levels were significantly increased after CRISPR/Cas9 RNP targeting of *PCSK1*-Prom, *PCSK1*-Enh and *PCSK1*- Prom+Enh, compared to the control (**Fig 4K-L**). Correspondingly, the average ratio of insulin secretion/ total proinsulin was reduced after targeting of *PCSK1* regulatory regions (**Fig S9D**). In sum, our studies with the CRISPR/Cas9 RNP approach suggest that the architecture and function of noncoding genomic regions can be interrogated in native human islet cells, including studies of gene regulation and β cell function after simultaneous targeting of distinct regulatory elements.

## DISCUSSION

Here we report successful genome editing in primary human islets using CRISPR/Cas9 RNP complexed with sgRNAs, revealing functions of *cis*-regulatory elements in human β cells. In the sole prior report of CRISPR/Cas9 targeting in primary human islets, we showed that coding and non-coding DNA in adult, post-mitotic pancreatic islet cells can be edited using lentivirus-based delivery of sgRNA and Cas9 enzyme^12^. However, lentiviral transduction of dispersed human islet cells in that study was relatively inefficient, precluding multiplex targeting or other studies. Electroporation of Cas9 RNP complexes has been used for genome editing in primary human T-cells and other dividing cell types^15–17^. However, adult human islet cells are post-mitotic, and it was unclear if Cas9 RNP delivery could achieve sufficient editing. Successful targeting of coding and non-coding DNA elements shown here should expand the uses of CRISPR for investigating human islet biology and islet-based diabetes risk.

Cas9 RNP complex electroporation efficiently targeted PDX1 coding sequence, and a postulated regulatory element previously linked by islet pc-HiC to *MPHOSPH9*, *PITPNM2* and *C12orf65*^8^. Our CRISPR/Cas9 RNP targeting showed selective impaired expression of the *MPHOSPH9* gene. *MPHOSPH9* encodes a protein required for cilia formation, an organelle essential for β and α cell function^22–23^. Moreover, reduced expression of *MPHOSPH9* was accompanied by decreased *INS* mRNA levels and insulin secretion. These findings highlight the potential for CRISPR/Cas9 RNP approaches to clarify the roles of cis-regulatory elements in islet gene regulation.

*PCSK1* is expressed at a high ’basal’ level in human β cells, and encodes the Prohormone Convertase 1/3 (PC1/3), an endopeptidase essential for Proinsulin processing. Glucose stimulates *PCSK1* expression, and prior studies demonstrated glucose-stimulated H3K27ac increases associated with the *PCSK1* promoter and a distal enhancer^8^, suggesting that these elements might regulate glucose-responsive *PCSK1* expression. In both T1D and T2D, PC1/3 activity appears reduced, and circulating Proinsulin:insulin ratios are increased^26^ (reviewed in^27^). These regulatory features motivated CRISPR targeting of these postulated *PCSK1 cis-*regulatory elements, including multiplexed targeting of promoter and enhancer. While we can not fully rule out the occurrences of large deletions in between the two targeted regulatory elements, two independent experiments indicate that such events might have occurred in <20% of the sequences. Our sequencing analysis did not reveal occurrence of duplication or reversion events. The use of a Cas9 RNP, that exhibits only short life and does not become integrated into the genome -as opposed to lentiCas9- might reduce the preponderance of such larger genomic rearrangements.

Multiplexed targeting with Cas9 RNP revealed that basal *PCSK1* expression was reduced only when both promoter and enhancer were mutated. This also resulted in a trend towards increased expression of α- cell markers, such as GCG and PCSK2, likely reflecting expected variation of islet transcriptomes between donors. By contrast, CRISPR/Cas9 RNP targeting revealed that each element is required for glucose-induced *PCSK1* expression, and impacts insulin secretion and total proinsulin levels. Overall, our approach revealed collaboration between *cis*-regulatory elements, a concept tested in other systems^16,28^, but not previously investigated by gene targeting in primary human islet cells. Further application of this CRISPR/Cas9 RNP approach could be fruitful in dissecting the role of elements containing additional candidate diabetes risk SNPs, including identification of their genetic targets in human islet cells. Together with prior work^12^, our results also show that efficiency of the targeting is locus dependent. CRISPR-based editing efficiency can be enhanced by targeting proliferating cells^12^; further studies are needed to assess if mitogenic stimulation of adult human islet cells, which are largely post-mitotic, could enhance CRISPR/RNP targeting efficiency. In addition, future efforts should also be directed to explore additional gene editing possibilities, such as engineered DNA-free virus-like particles (VLPs)^32^, base editing^33^ and prime editing^34^ strategies. One caveat of our studies is that we were unable to correlate phenotypes in single cells or clonal populations with gene editing. The non-dividing nature of islet cells precludes the enrichment studies usually performed for cell lines. Approaches such as TARGET-seq^35^ or other single cell-based approaches could be useful in future studies to assess gene-editing heterogeneity.

In summary, CRISPR/Cas9 RNP electroporation in primary human islets allowed: (1) highly efficient targeting (deletion) of non-coding DNAs, (2) simultaneous deletion of two regulatory elements, and (3) functional assessment of regulatory impacts from multiplex targeting of non-coding DNA, including dynamic physiological regulation by glucose. These innovations advance the use of genome editing to dissect the genetic regulatory mechanisms in islets that underlie diabetes risk.

## Acknowledgments

We thank past and current members of the Kim group, especially S. Park, for technical advice and support. We thank Dr. Jocelyn Manning Fox, Mrs. Nancy Smith, and Mr. James Lyon (University of Alberta) for their contributions to human islet isolations. We gratefully acknowledge organ donors and their families, Canadian organ procurement organizations, particularly the Human Organ Procurement and Exchange (HOPE) program and the Trillium Gift of Life Network, and islet procurement through the Alberta Diabetes Institute Islet Core, Integrated Islet Distribution Program (U.S. NIH UC4 DK098085), the National Disease Research Interchange, and the International Institute for the Advancement of Medicine. R.J.B. was supported by a postdoctoral fellowship from JDRF (3-PDF-2018-584- A-N). ALG is supported by a Wellcome Senior Fellowship (200837/Z/16/Z) and by NIDDK UM-1DK126185. Work in the Kim lab was supported by NIH awards (R01 DK107507; R01 DK108817; U01 DK123743 to S.K.K.), and JDRF Northern California Center of Excellence (to S.K.K. and A.M.). Work here was also supported by NIH grant P30 DK116074 (S.K.K.), and by the Stanford Islet Research Core, and Diabetes Genomics and Analysis Core of the Stanford Diabetes Research Center.

## Author contributions

R.J.B. and S.K.K. conceptualized the study and guided the work. Methodology: R.J.B., A.L.G., S.K.K.; Investigation: R.J.B., S.H.K., W.Z., A.M.; Writing: R.J.B. and S.K.K. wrote the manuscript with input from all coauthors; Visualization: R.J.B.; Supervision: S.K.K.; Funding acquisition: R.J.B., A.M., A.L.G., S.K.K.

## Declaration of interests

R.J.B., S.H.K., W.Z., and S.K.K. declare no competing interests. A.M. is a co-founder of Arsenal Biosciences, Spotlight Therapeutics, and Survey Genomics, serves on the boards of directors at Spotlight Therapeutics and Survey Genomics, is a board observer (and former member of the board of directors) at Arsenal Biosciences, is a member of the scientific advisory boards of Arsenal Biosciences, Spotlight Therapeutics, Survey Genomics, NewLimit, Amgen, Tenaya, and Lightcast. A.M. owns stock in Arsenal Biosciences, Spotlight Therapeutics, NewLimit, Survey Genomics, PACT Pharma, Tenaya, and Lightcast and has received fees from Arsenal Biosciences, Spotlight Therapeutics, NewLimit, Survey Genomics, 23andMe, PACT Pharma, Juno Therapeutics, Tenaya, Lightcast, Trizell, Vertex, Merck, Amgen, Genentech, AlphaSights, Rupert Case Management, Bernstein, and ALDA. A.M. is an investor in and informal advisor to Offline Ventures and a client of EPIQ. The Marson laboratory has received research support from Juno Therapeutics, Epinomics, Sanofi, GlaxoSmithKline, Gilead, and Anthem. A.L.G.’s spouse holds stock options in Roche and is an employee of Genentech.

**Supplemental Figure 1. Efficient CRISPR/Cas9 RNP mediated targeting of PDX1 in primary human islet cells.** (A) Quantification of indels in the *PDX1* exon sequence after electroporation of CRISPR/Cas9 RNP complexed with PDX1 sgRNAs, comparing 2 different concentrations of CRISPR/Cas9 RNP complexes (1X and 2X) and using 3 different voltages (V1-V3) (see methods). (B) % of live cells after electroporation with Cas9 RNP +sgRNAs relative to the non-electroporated control (n=2). (C-E) FlowJo Plots following Live/Dead staining of human pseudoislets: (B) unstained control, (C) Live/Dead non-electroporated group, (D) Live/Dead Cas9 RNP-2sgRNAs complex electroporated islet cells. (F) Alleles frequency table around PDX1 sgRNA, generated with CRISPResso. The predicted cleavage position is shown as a vertical dotted line, insertions as shown in a red box, deletions as a dotted line.

**Supplemental Figure 2.** pcHi-C chromatin looping in the *MPHOSPH9* locus. Data obtained from the IsletRegulomeBrowser^29^.

**Supplemental Figure 3. Bioinformatics assessment of potential off-target genomic sites.** (A) Potential off-target sites were identified using Chop-Chop^30^. (B) TIDE analysis showing that no indels were detected in the off-target sites. (C) PCR-amplification followed by sequencing of predicted off-target sites for the sgRNAs specific to the presumptive enhancer in the MPHOSPH9 locus. For the 3 potential off-targets sites evaluated for each sgRNA used (MPHOSPH9_Enhancer_sg1F and MPHOSPH9_Enhancer_sg1R), only wild-type sequences were detected. Red lines highlight the potential off-target sgRNA sequence. * indicates mismatched nucleotides. This analysis was repeated for 3 independent CRISPR-Targeted donors.

**Supplemental Figure 4. PCSK1 regulation in human islets.** (A) Islet Regulome Browser tracks of *PCSK1*. Virtual 4C representations show medium-confidence interactions (CHICAGO score 3-5, blue) between *PCSK1*-Enh element (pink box) and *PCSK1*-Prom (turquoise box) in human islets. The HindIII fragment that contains *PCSK1*-Prom is used as viewpoint and depicted as inverted triangles. Diagram candidate T2D risk SNPs overlapping the Enhancer region are shown. Open chromatin classes in the *PCSK1* locus in human islets include putative active enhancer (red), active promoter (green), inactive enhancer (grey) and CTCF (blue).

**Supplemental Figure 5. CRISPR/Cas9 RNP electroporation in human islet cells allows multiplexed targeting of regulatory regions.** (A,E) Alleles frequency tables generated with CRISPResso around (A) *PCSK1*-Prom and (E) *PCSK1*-Enh deleted regions, after multiplexed targeting of both *PCSK1*-Prom and *PCSK1*-Enh regions. (B) Histogram of Summary of Promoter deletions within the region targeted by *PCSK1*-Prom sgRNA1 and sgRNA2, (C-D) alleles frequency table with the viewpoint in (C) *PCSK1*-Prom sgRNA1 or (D) *PCSK1*-Prom sgRNA2. (F) Histogram of Summary of Enhancer deletions within the region targeted by *PCSK1*-Enh sgRNA1 and sgRNA2, (G-H) alleles frequency table with the viewpoint in (G) *PCSK1*-Enh sgRNA1 or (H) *PCSK1*-Enh sgRNA2. The predicted cleavage position is shown as a vertical dotted line, deletions as a dotted line.

**Supplemental Figure 6. Bioinformatics assessment of potential off-target genomic sites induced by CRISPR/Cas9 and sgRNAs targeting a presumptive enhancer and/or promoter in the PCSK1 locus**: Potential off-target sites were identified using Chop-Chop^30^. PCR-amplification followed by sequencing of predicted off-target sites for *PCSK1*_Enhancer_sg1R and *PCSK1*_Enhancer_sg1F. For the 3 potential off-target sequences tested for each sgRNA, wild-type sequences were detected. The sgRNAs designed to target the promoter sequence specific retrieved no potential off-targets. Red boxes highlight the presumptive off-target sequences. This analysis was repeated for 3 independent donors and all the conditions *PCSK1*-Prom, *PCSK1*-Enh and *PCSK1*-Enh+Prom.

**Supplemental Figure 7.** Measurement of proportion of internal amplicons to external reference amplicon following CRISPR targeting with 4 sgRNAs in the *PCSK1* locus. (A) Design of indexed PCR: three internal PCRs (int1, int2 and int3) were performed in the 28 kB region in between targeted *PCSK1* enhancer and Promoter. A third PCR was designed outside the targeted region (ref). (B) Following Miseq sequencing, amplicons were quantified with CRISPResso. The ratios of internal/reference amplicons for 3 independent amplicons were calculated for *PCSK1*-Prom+Enh (n= 3 independent donors). (C-D) Measurements were also included for PCR int1 (C) and for PCR int2 (D), for *PCSK1*-Prom and *PCSK1*-Enh.

**Supplemental Figure 8.** Assessment of Proportion of Large deletions following CRISPR targeting with 4 sgRNAs in the *PCSK1* locus. (A) Design of multiplexed PCR: PCR was performed with two primer pairs, one external to sgRNAs targeting *PCSK1*-Prom and *PCSK1*-Enh to detect large deletions (LDs, P_1F and E_1R), and a second, internal one, to detect the wild-type sequence in between *PCSK1*-Prom and *PCSK1*-Enh (Pint_1F and Pint_1R). Large deletions can be consequence of cutting by *PCSK1*_Prom_sg1 or sg2 and *PCSK1*_Enh sg3 or sg4, resulting in two different large deletion products (LD1/4 and LD2/3). (B) Agarose gel showing multiplexed PCR for samples: lane 1-Cas9-NT control, lane 2- *PCSK1*_Prom_donor1 (P_P1), lane 3-*PCSK1*_Prom+Enh_donor 1 (P_P+E1), lane 4- *PCSK1*_Prom+Enh donor 2 (P_P+E2), lane 5-*PCSK1*_Prom_donor 2 (P_P2), lane 6- *PCSK1*_Enh_donor 2 (P_E2) and *PCSK1*_Prom+Enh donor 3 (P_P+E3). (C-D) Quantification of proportion of large deletion bands (LD1 and LD2), normalized to amplicon size, after background subtraction (see Methods).

**Supplemental Figure 9.** (A-B) RT-qPCR after CRISPR/Cas9 RNP targeting of *PCSK1*-Prom (turquoise), *PCSK1*-Enh (pink) or *PCSK1*-Enh+Prom (green): (A-B) Following culture at 5.6 mM glucose, and measurement of mRNA levels of (A) *INS* and (B) *IAPP* (n = 2 donors) relative to the 2.8 mM cultured control. (C) Expression level of PCSK2 and GCG following culture at 16.7 mM relative to 2.8 mM glucose (n = 2 independent donors). (D) Measurements of the ratio of insulin secretion/ total proinsulin content for Cas9-sgNT, *PCSK1*-Prom, *PCSK1*-Enh and *PCSK1*-Enh+Prom (3 independent donors).

## STAR Methods

### Human Islet Procurement

Organs and islets were procured through the Integrated Islet Distribution Network (IIDP), National Diabetes Research Institute (NDRI) and the Alberta Diabetes Institute (ADI) Islet Core. De-identified human islets were obtained from healthy, non-diabetic organ donors with less than 18 h of cold ischemia time, and deceased due to acute trauma or anoxia. For this study, islets from 22 adult donors were used (Supplementary Table 1).

### CRISPR/Cas9 RNP complexes assembly

Cas9 from Streptococcus pyogenes, fused with enhanced GFP, recombinant, expressed in E. coli, 3X NLS was purchased from Sigma-Aldrich (CAS9GFPPRO). The sgRNAs used in this study were designed using E-CRISPR and MIT CRISPR design tool. Chemically modified sgRNAs (with 2’-O-Methyl at 3 first and last bases, and 3’ phosphorothiate bonds between first 3 and last 2 bases) were purchased from Synthego. For CRISPR/Cas9 RNP complexes assembly, Cas9GFPPRO was complexed to sgRNAs and incubated at room temperature for 15 min. For 1X concentration, 1250 ng Cas9GFPPRO were incubated with 7.5 pg sgRNAs in 5 ul buffer R (NeonTM, Thermo Fisher Scientific). For 2X condition, double concentration was used (Supplemental Figure 1A). The sgRNA sequences used in this study can be found in Supplementary Table 2.

Human pseudoislet generation and CRISPR/Cas9 RNP complexes electroporation Human islets were dispersed into a single cell suspension by enzymatic digestion (Accumax, Invitrogen). For each electroporation pulse with 10 ul tips of the NeonTM transfection system (Thermo Fisher Scientific), 150,000 cells were resuspended in 5 ul buffer R. Following CRISPR/Cas9 RNP assembly, CRISPR/Cas9 RNP complexes were added to cells in buffer R. Electroporation conditions tested in Supplemental Fig. 1A consisted of: V1 = 1100V, 1 pulse, 40 ms, V2 = 1500 V, 3 pulses, 10 ms, V3 = 1600 V, 3 pulses, 10 ms. V3 was used for following experiments. Immediately after the pulse, cells were transferred into 24 well plates in culture medium comprised of RPMI 1640 (Gibco), 2.25 g/dl glucose, 1% penicillin/streptomycin (v/v, Gibco), and 2% human serum (Sigma-Aldrich).

Electroporated islet cells were cultured in an Orbi-Shaker (34-206, Genesee Scientific) until day 5 or 6 prior to further molecular or physiological analysis. One day following electroporation, GFP expression was evaluated.

### Live/Dead staining and flow cytometry of human islet cells

Pseudoislets were dispersed into single cells -as detailed before-, resuspended in PBS and stained with LIVE/DEAD Fixable Near-IR dead cell stain kit (Life Technologies) for 30 min in the dark. A negative, unstained control was included. Labeled cells were run on the Aurora 2 (Cytek Biosciences) using appropriate compensation controls and doublet removal. Quantification of Live/Dead cells was performed using FlowJo.

### Genomic DNA extraction, MiSeq sequencing, and CRISPResso analysis

1000–5000 electroporated cells were used for genomic DNA (gDNA) extraction using the Arcturus® PicoPure® DNA Extraction Kit (Thermo Fisher Scientific) per the manufacturer’s instructions. Next, extracted gDNA was amplified using the REPLI-g-mini Kit (Qiagen). 2 ul of extracted and amplified gDNA was used for nested PCR using Phusion U Green Multiplex PCR Master Mix (2X) (Thermo Scientific). PCR1 was performed with amplicon specific and barcode tail primers, with conditions consisting of initial 98 °C for 2 min, followed by 30 cycles, each composed of a denaturing step at 98 °C for 10 s, an annealing step at 61 °C for 20 s, and an extension step at 72 °C for 30 s, followed by a final extension at 72 °C for 2 min. PCR2 was included to add barcodes for sequencing, using publicly available Illumina Adapter sequences. Conditions consisted of initial 98 °C for 2 min, followed by 12 cycles, each composed of a denaturing step at 98 °C for 10 s, an annealing step at 61 °C for 20 s, and an extension step at 72 °C for 30 s, followed by a final extension at 72 °C for 2 min. PCR2 products were gel purified, using Zymoclean Gel DNA recovery kit (Zymo Research). Amplicon concentrations were measured using Qubit dsDNA HS (Thermo Fisher Scientific) and pooled for Miseq sequencing at 2x300 bp (Illumina, Stanford Functional Genomics Facility). CRISPResso analysis was performed of the sequenced amplicons, following the software pipeline designed by the Pinello Lab^20^. Primer sequences are summarized in Supplementary Table 3.

### Evaluation of CRISPR/Cas9 off-target effects and PCRs to detect large deletions

Off-target site prediction was performed using the CHOP-CHOP tool^30^. The potential off-target sites found had at least 2 mismatches with respect to the sgRNA sequence. PCR primers encompassing 3 of these potential off-target sites for each sgRNA were designed and the PCR amplicons were purified and sequenced. PCR was performed with Phusion U Green Multiplex PCR Master Mix (2X) (Thermo Scientific) in 35 cycles, each cycle composed of a denaturing step at 95 °C for 1 min, an annealing step at 61 °C for 30 sec and an extension step at 68 °C for 1 min, followed by final extension at 72°C for 5 min. Primer sequences are summarized in Supplementary Table 3. Following sequencing, we performed TIDE^31^ analyses to assess indel occurrence at these off target sites.

To assess occurrence of deletions of the region in between the targeted PCSK1 enhancer and promoter, we performed nested indexed amplification of 4 amplicons (three within the potentially deleted region -intPCR1, intPCR2 and intPCR3- and one reference amplicon, located outside the presumably deleted region, **Fig. S7A**). PCR1 and PCR2 were performed with Phusion U Green Multiplex PCR Master Mix (2X) (Thermo Scientific), gel purified, Miseq sequenced and CRISPResso analyzed as described above. The proportion of internal PCR amplicons/reference amplicon were calculated both for the control and *PCSK1*_Prom+Enh targeted groups. Primer sequences are included in Supplementary Table 3. To directly measure amplicons generated by deletion of the entire region in between the targeted promoter and enhancer, we performed multiplexed PCR, design shown in **Fig. S8A**. Multiplex PCR was performed with Phusion U Green Multiplex PCR Master Mix (2X) (Thermo Scientific) in 30 cycles, each cycle composed of a denaturing step at 95 °C for 1 min, an annealing step at 65 °C for 20 sec and an extension step at 72 °C for 15 sec, followed by a final extension at 72°C for 5 min. Primer sequences are summarized in Supplementary Table 3. PCR products were run on a 1.5% low melt agarose gel for 40 minutes under a fixed voltage of 110 Volts. The gel was imaged using Axygen, gel documentation systems (Corning). Quantitation was performed via FIJI (ImageJ). A selection box with a fixed area was generated and utilized to report the mean intensity and area under the curve for all PCR products per condition. The background was subtracted for each lane, and the quantitation was normalized according to the product’s size (bp).

### RNA extraction and quantitative RT-PCR

RNA was isolated from electroporated pseudoislet cells using the PicoPure RNA Isolation Kit (Life Technologies). cDNA was synthesized using the Maxima First Strand cDNA synthesis kit (Thermo Scientific) and gene expression was assessed by PCR using the Taqman Gene Expression Mix (Thermo Scientific). Data were analyzed using Prism 6.0 h (GraphPad Software Inc., San Diego, CA). Taqman probes used for this study are summarized in Supplementary Table 3. Paired two-tailed t tests were used to indicate statistical significance, and data are presented as mean and standard deviation.

### Glucose-Stimulated human insulin and proinsulin secretion, insulin, and proinsulin content measurement

Glucose-Stimulated human insulin and proinsulin secretion were performed as batch assays on pseudoislets from CRISPR-Control and Cas9 targeted groups, with 30 pseudoislets as input, and supernatants were collected after 1 h incubation at 2.8 mM, 16.7 mM and 16.7 mM +IBMX glucose concentrations. To determine total cellular insulin or proinsulin content, pseudoislets were sonicated and lysed to extract the total cellular insulin or proinsulin content (Human insulin and proinsulin ELISA kits, Mercodia). gDNA from the same sonicated islets used for the insulin ELISAs was extracted and used for normalization. Data were analyzed using Prism 6.0 h (GraphPad Software Inc., San Diego, CA), normalized to the CRISPR-Control and presented as mean with standard deviation. Paired two-tailed t tests were used to indicate statistical significance.

### Data visualization

Browser tracks were made with the UCSC genome browser. The graphics were made with BioRender.

