## Supplemental Fig 1-9 for "Multiplexed CRISPR gene editing in primary human islet cells with Cas9 ribonucleoprotein"

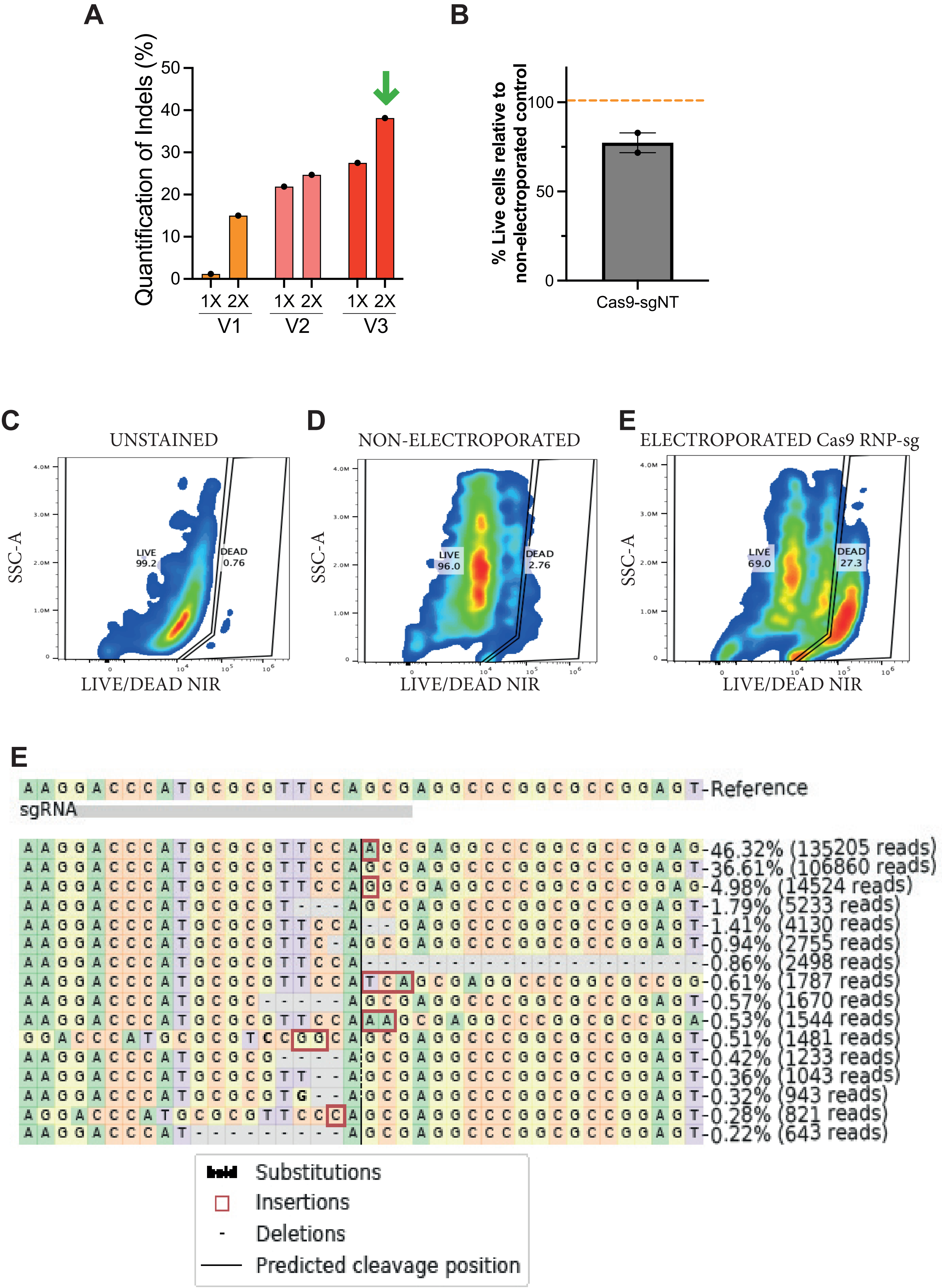

A
Quantification of Indels (%)
0
10
20
30
40
50
1X 2X
V1
1X 2X
V2
1X 2X
V3
B
% Live cells relative to non-electroporated control
0
50
100
Cas9-sgNT
C
UNSTAINED
SSC-A
LIVE 99.2
DEAD 0.76
LIVE/DEAD NIR
D
NON-ELECTROPORATED
SSC-A
LIVE 96.0
DEAD 2.76
LIVE/DEAD NIR
E
ELECTROPORATED Cas9 RNP-sg
SSC-A
LIVE 69.0
DEAD 27.3
LIVE/DEAD NIR
E
AAGGACCCATGCGCGTTCCAGCGAGGCCCGGCGCCGGAGT-Reference
sgRNA
-46.32% (135205 reads)
-36.61% (106860 reads)
-4.98% (14524 reads)
-1.79% (5233 reads)
-1.41% (4130 reads)
-0.94% (2755 reads)
-0.86% (2498 reads)
-0.61% (1787 reads)
-0.57% (1670 reads)
-0.53% (1544 reads)
-0.51% (1481 reads)
-0.42% (1233 reads)
-0.36% (1043 reads)
-0.32% (943 reads)
-0.28% (821 reads)
-0.22% (643 reads)
Substitutions
Insertions
Deletions
Predicted cleavage position

A

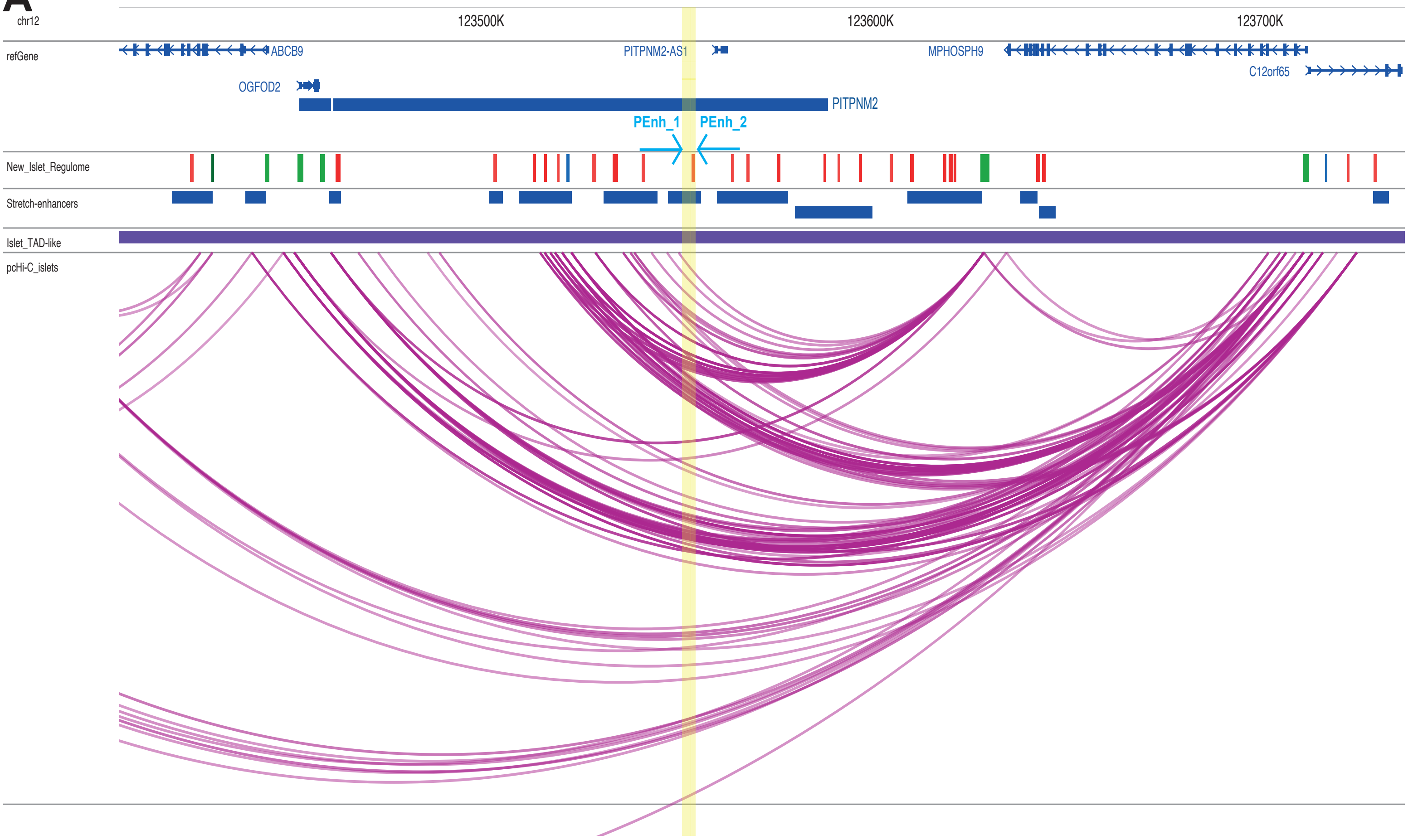
chr12
123500K
123600K
123700K
refGene
ABCB9
PITPNM2-AS1
MPHOSPH9
C12orf65
OGFOD2
PITPNM2
PEnh_1
PEnh_2
New_Islet_Regulome
Stretch-enhancers
Islet_TAD-like
pcHi-C_islets

**A** Off targets

**MPHOSPH9_1F**

| Location | No. mmatches | Sequence (mismatches in red) |
|---|---|---|
| chr3:28616370 | 3 | aAGTGGTGACaTcACTTGAAAGG |
| chr6:125927687 | 2 | CCTTTCAAGTTcAGTgACCACTA |

**MPHOSPH9_1R**

| Location | No. mmatches | Sequence (mismatches in red) |
|---|---|---|
| chr14:25603174 | 3 | CCTGAGGTAGATtGTGtGTGCTt |
| chr6:19720047 | 3 | GAGCACgCAgGATCTACCTtGGG |
| chr6:44064245 | 3 | CCCGAGGaAGAcCGTGAGTGCcC |
| chr8:41615203 | 3 | CCGGAGGagGATgGTGAGTGCTC |

**B**

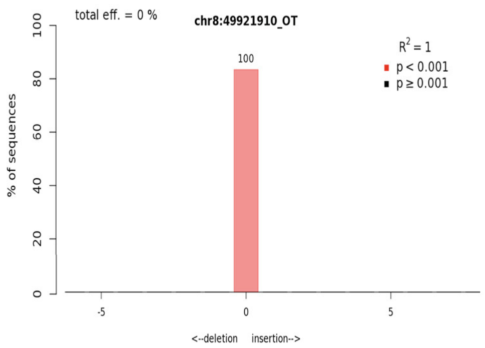


**C**

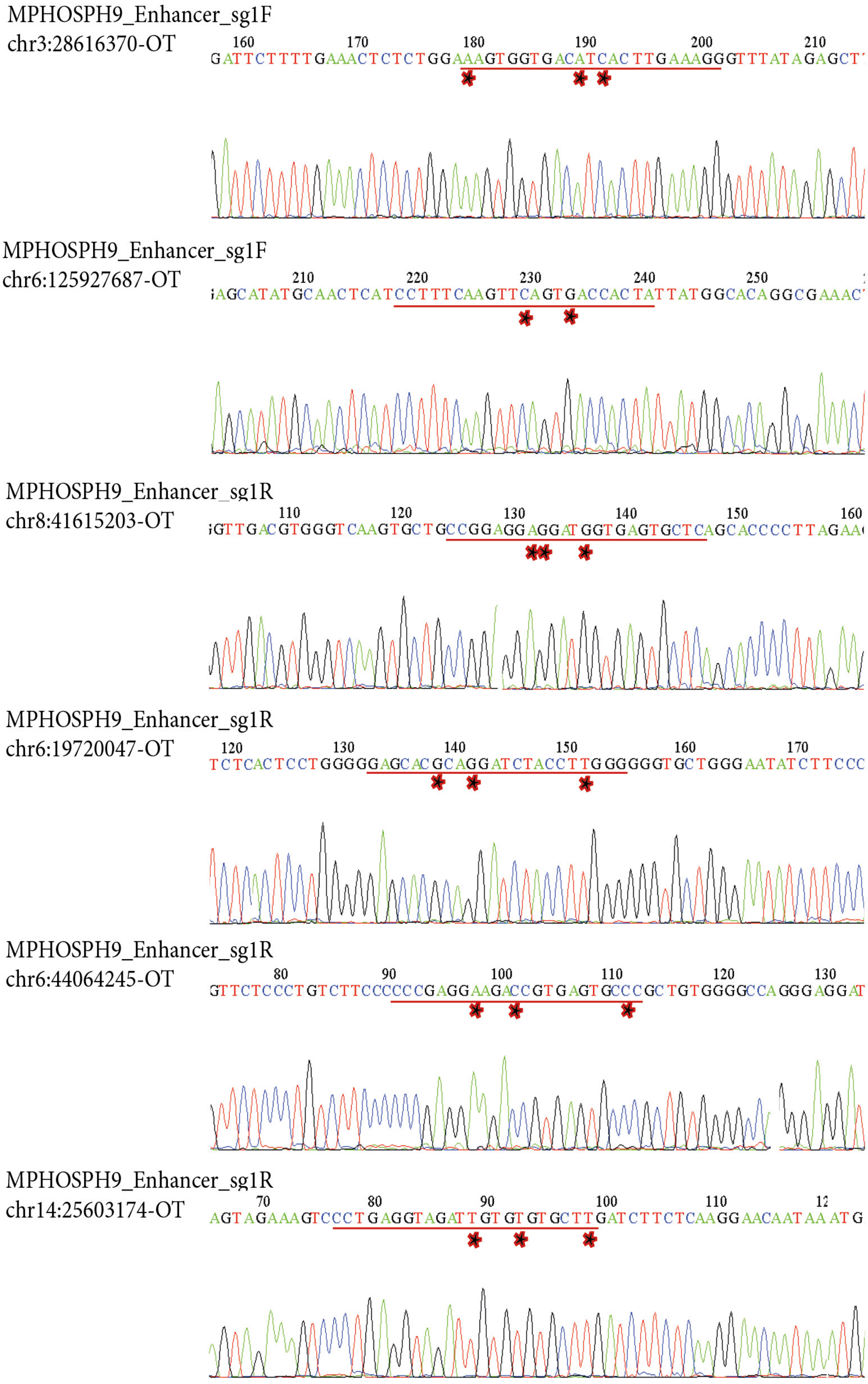

A

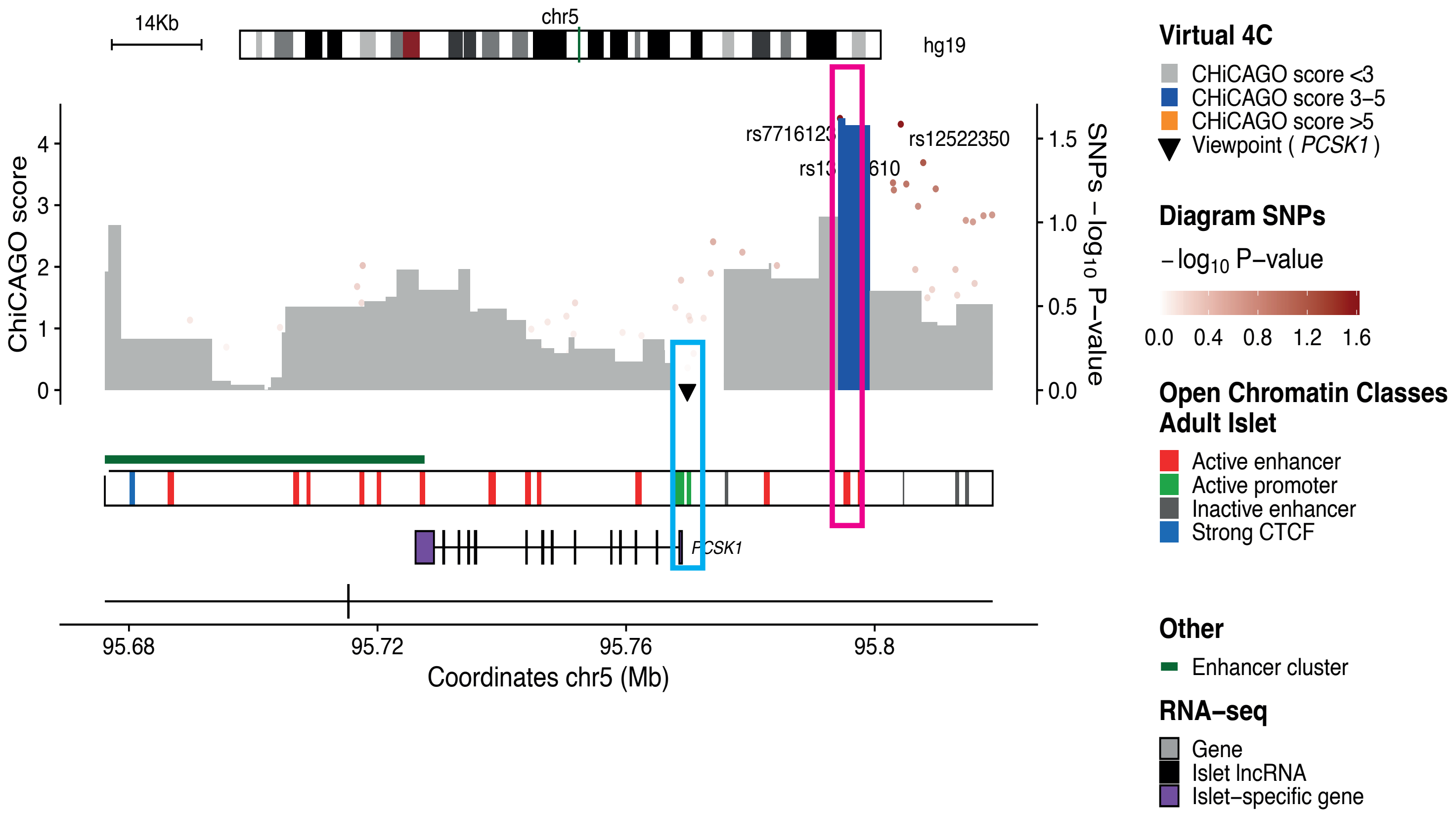
14Kb
chr5
hg19
ChiCAGO score
SNPs – log10 P–value
rs7716123
rs12522350
PCSK1
Coordinates chr5 (Mb)
95.68
95.72
95.76
95.8
Virtual 4C
CHiCAGO score <3
CHiCAGO score 3–5
CHiCAGO score >5
Viewpoint ( PCSK1 )
Diagram SNPs
– log10 P–value
0.0 0.4 0.8 1.2 1.6
Open Chromatin Classes
Adult Islet
Active enhancer
Active promoter
Inactive enhancer
Strong CTCF
Other
Enhancer cluster
RNA–seq
Gene
Islet lncRNA
Islet–specific gene

### PCSK1_Enhancer+Promoter targeting

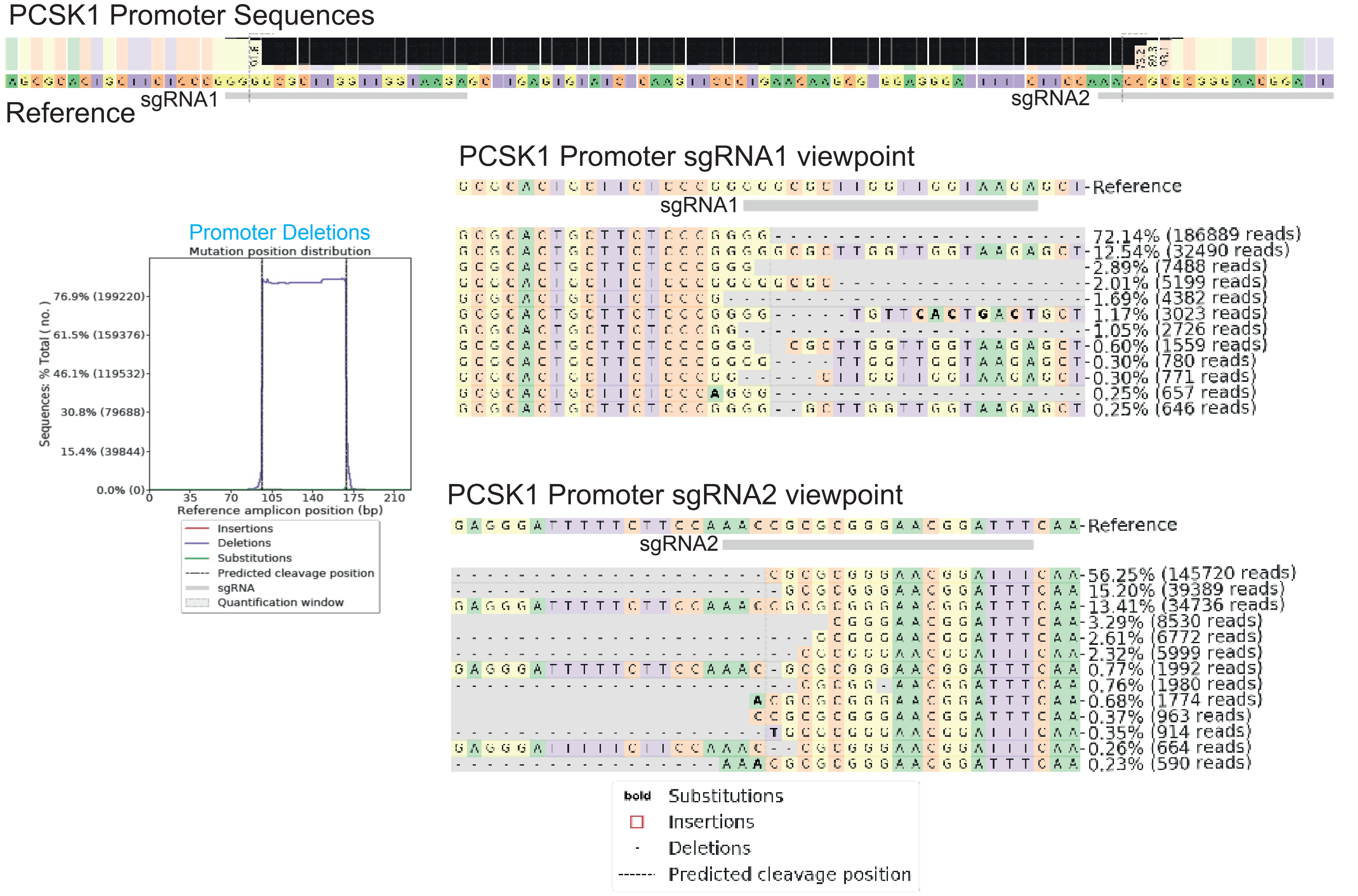


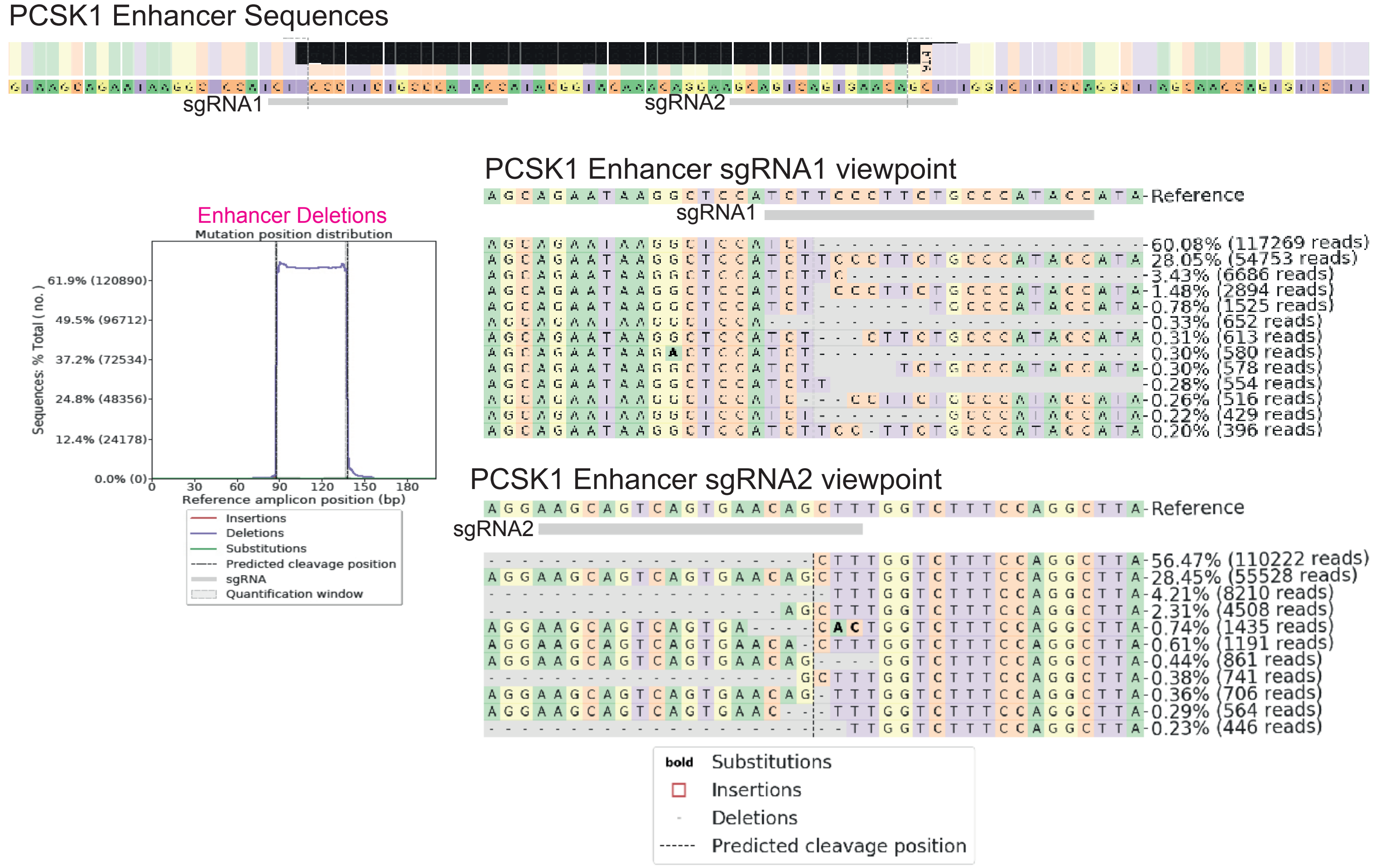

PCSK1_Enhancer_sg1R
chr1:161952532-OT

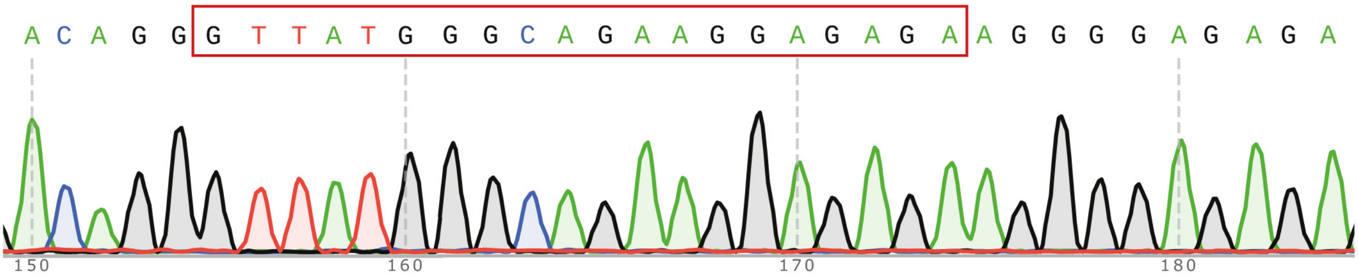


PCSK1_Enhancer_sg1R
chr18:32365907-OT

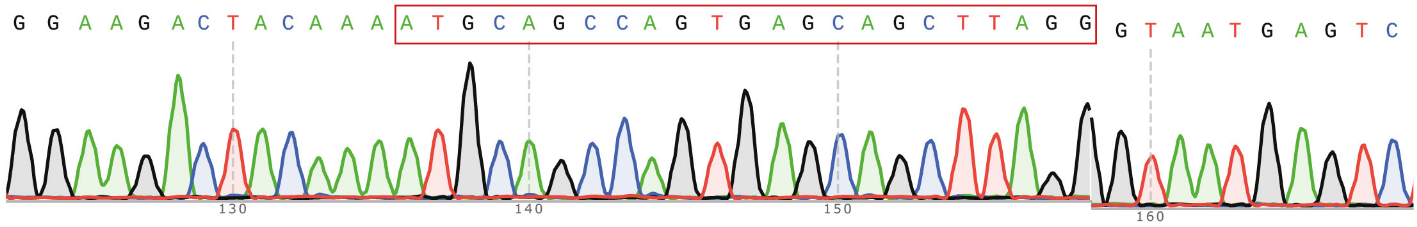


PCSK1_Enhancer_sg1R
chr10:93487121-OT

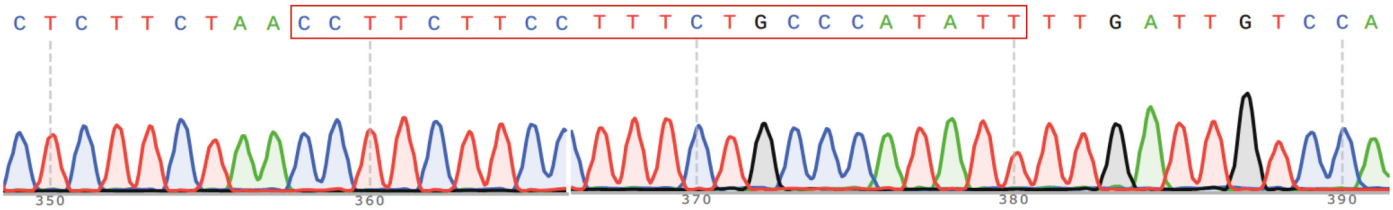


PCSK1_Enhancer_sg1F
chr11:56543423-OT

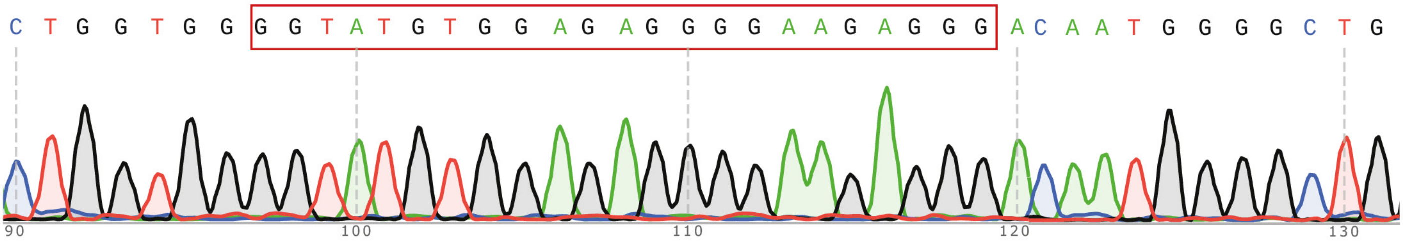


PCSK1_Enhancer_sg1F
chr13:95247995-OT

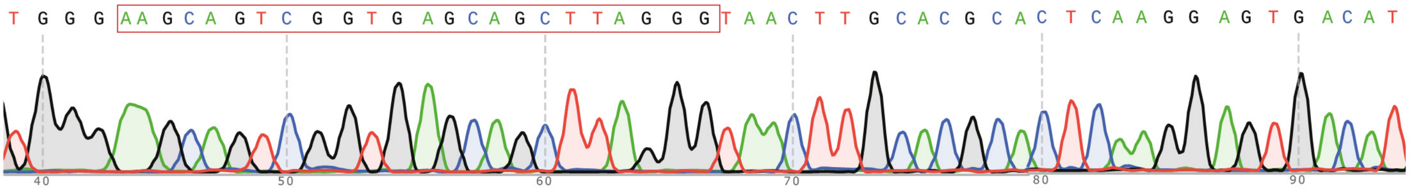


PCSK1_Enhancer_sg1F
chr3:59872738-OT

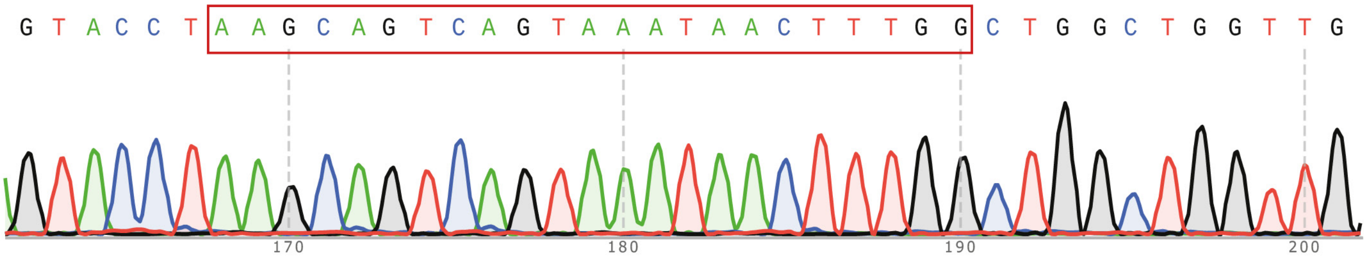

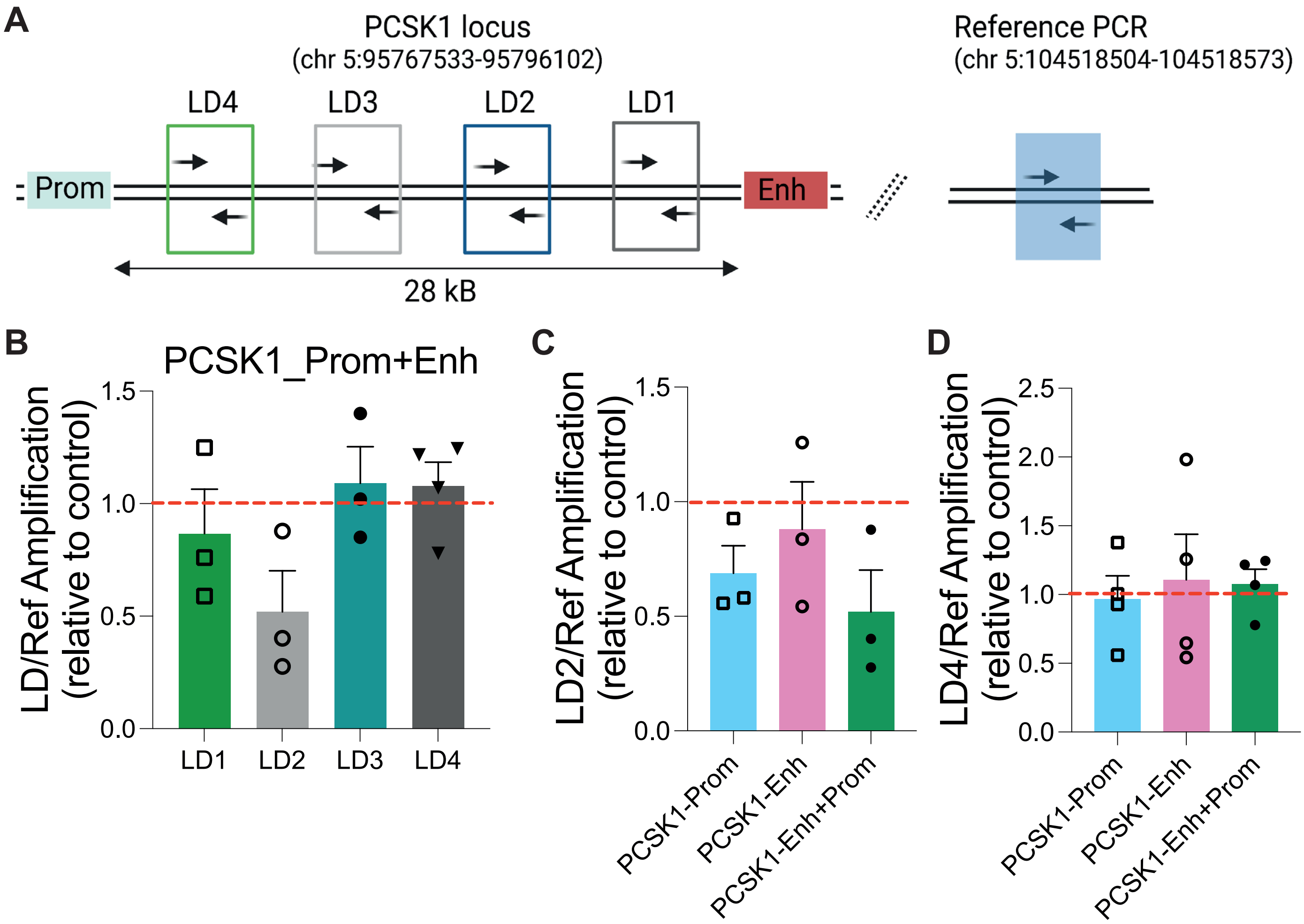
A
PCSK1 locus
(chr 5:95767533-95796102)
LD4
LD3
LD2
LD1
Prom
Enh
28 kB
Reference PCR
(chr 5:104518504-104518573)
B
PCSK1_Prom+Enh
LD/Ref Amplification (relative to control)
1.5
1.0
0.5
0.0
LD1
LD2
LD3
LD4
C
LD2/Ref Amplification (relative to control)
1.5
1.0
0.5
0.0
PCSK1-Prom
PCSK1-Enh
PCSK1-Enh+Prom
D
LD4/Ref Amplification (relative to control)
2.5
2.0
1.5
1.0
0.5
0.0
PCSK1-Prom
PCSK1-Enh
PCSK1-Enh+Prom

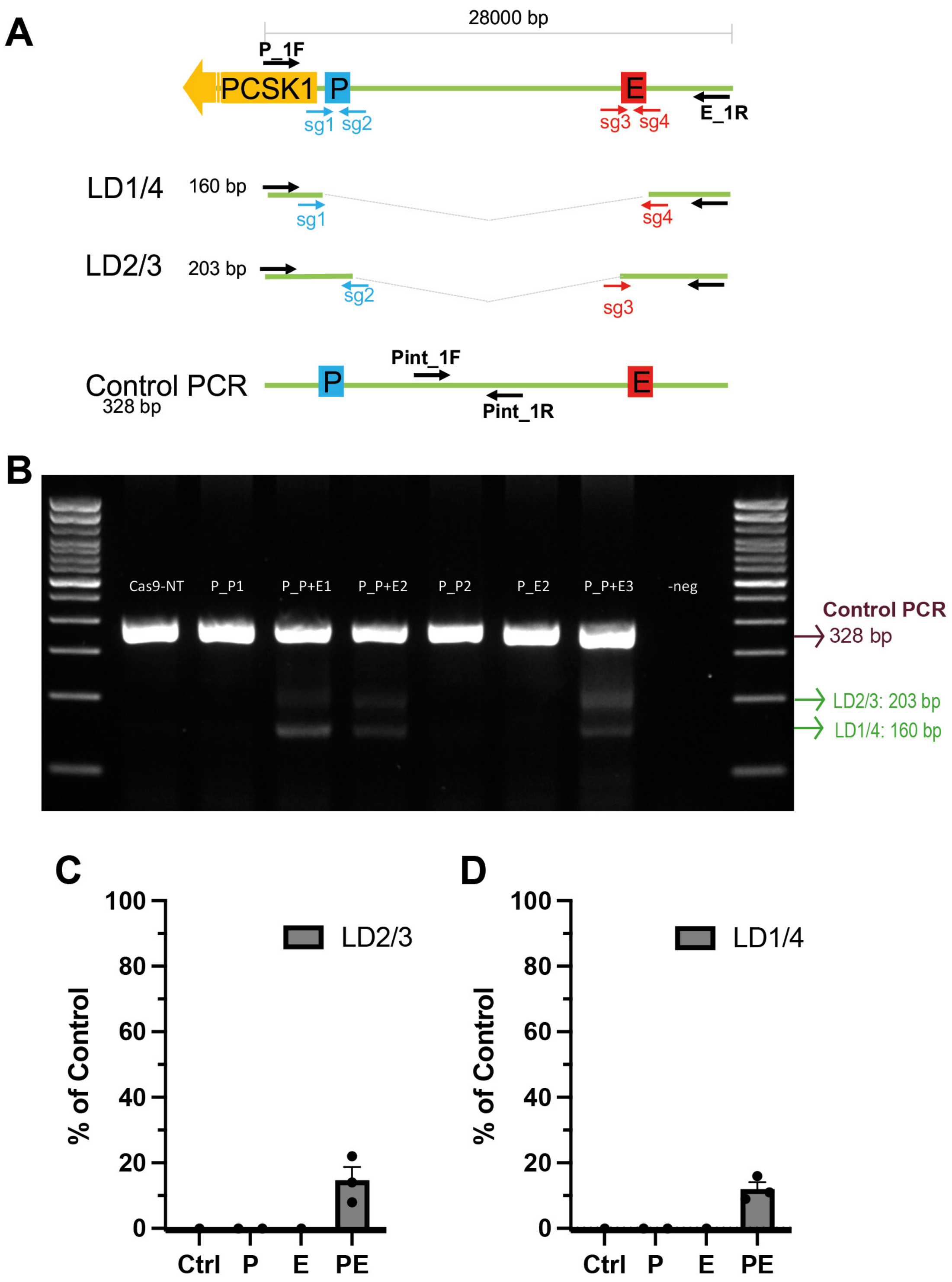
A
28000 bp
P_1F
PCSK1
P
E
sg1 sg2
sg3 sg4
E_1R
LD1/4
160 bp
sg1
sg4
LD2/3
203 bp
sg2
sg3
Control PCR
328 bp
Pint_1F
Pint_1R
B
Cas9-NT
P_P1
P_P+E1
P_P+E2
P_P2
P_E2
P_P+E3
-neg
Control PCR
328 bp
LD2/3: 203 bp
LD1/4: 160 bp
C
% of Control
LD2/3
0
20
40
60
80
100
Ctrl
P
E
PE
D
% of Control
LD1/4
0
20
40
60
80
100
Ctrl
P
E
PE

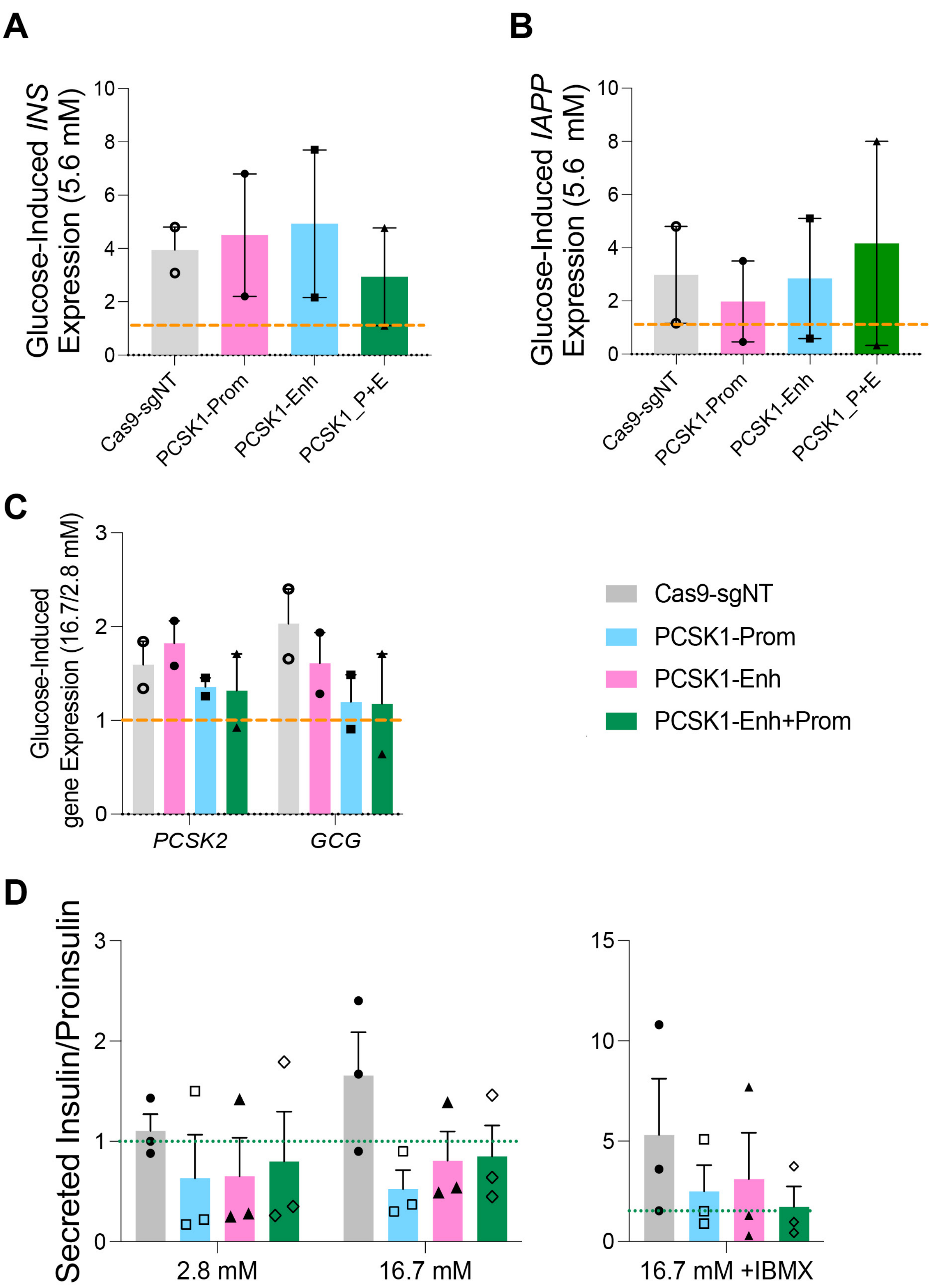
A
Glucose-Induced INS Expression (5.6 mM)
10
8
6
4
2
0
Cas9-sgNT
PCSK1-Prom
PCSK1-Enh
PCSK1_P+E
B
Glucose-Induced IAPP Expression (5.6 mM)
10
8
6
4
2
0
Cas9-sgNT
PCSK1-Prom
PCSK1-Enh
PCSK1_P+E
C
Glucose-Induced gene Expression (16.7/2.8 mM)
3
2
1
0
PCSK2
GCG
Cas9-sgNT
PCSK1-Prom
PCSK1-Enh
PCSK1-Enh+Prom
D
Secreted Insulin/Proinsulin
3
2
1
0
2.8 mM
16.7 mM
15
10
5
0
16.7 mM +IBMX
